# Seascape genomics reveals shared spatial genetic structure in three co-occurring sea snakes

**DOI:** 10.64898/2026.10.02.756145

**Authors:** Shannon Coppersmith, Vinay Udyawer, Jenna Margaret Crowe-Riddell, James Henry Nankivell, Amelia Rose Pointon, Vhon Oliver Garcia, Blanche d’Anastasi, Claire Goiran, Kate Laura Sanders, Alastair John Ludington

## Abstract

Marine species with broad distributions often exhibit complex patterns of genetic connectivity, shaped by historical and contemporary processes. Understanding these patterns is critical for conserving taxa threatened by ongoing and future environmental change. Here, we use reduced-representation and whole-genome sequencing to investigate population genetic patterns in three co-occurring sea snake species (*Hydrophis stokesii*, *Hydrophis major*, and *Aipysurus laevis*) across northern Australia and New Caledonia. Broadscale analyses revealed major genetic breaks across all species between: (1) Australia and New Caledonia, (2) northwestern and northeastern Australia, and (3) the offshore Timor Sea reefs and adjacent northwestern continental shelf of Australia. More subtle genetic differentiation was observed between Shark Bay in Western Australia and northern localities. All species showed relatively large effective population sizes and broadscale connectivity across ∼1,300 km of Australia’s northwestern continental shelf. This likely reflects substantial historical range expansion over late Quaternary timescales, which may have contributed to the observed genetic connectivity across the region. However, subtle patterns of isolation by distance emphasised the contribution of species-specific traits in shaping connectivity. The isolated offshore reef populations are genetically distinct and potentially threatened by environmental change. Disjunct populations in Shark Bay and the Timor Sea maintain similar levels of genetic diversity to the extensive continental shelf populations, while New Caledonian populations exhibit reduced diversity. Our results support the delineation of spatial units for sea snake conservation based on shared phylogeographic structure, while also finding important population- and species-specific differences in connectivity at finer spatial and temporal scales.

**Highlights:**

- Broadscale phylogeographic structure is shared by three species of sea snakes.
- Populations along ∼1,300 km of the NW Australian shelf show extensive genetic connectivity.
- Fine-scale patterns highlight the role of species traits in shaping connectivity.
- Conservation planning must consider both broadscale and species-specific processes.

**Graphical Abstract:** 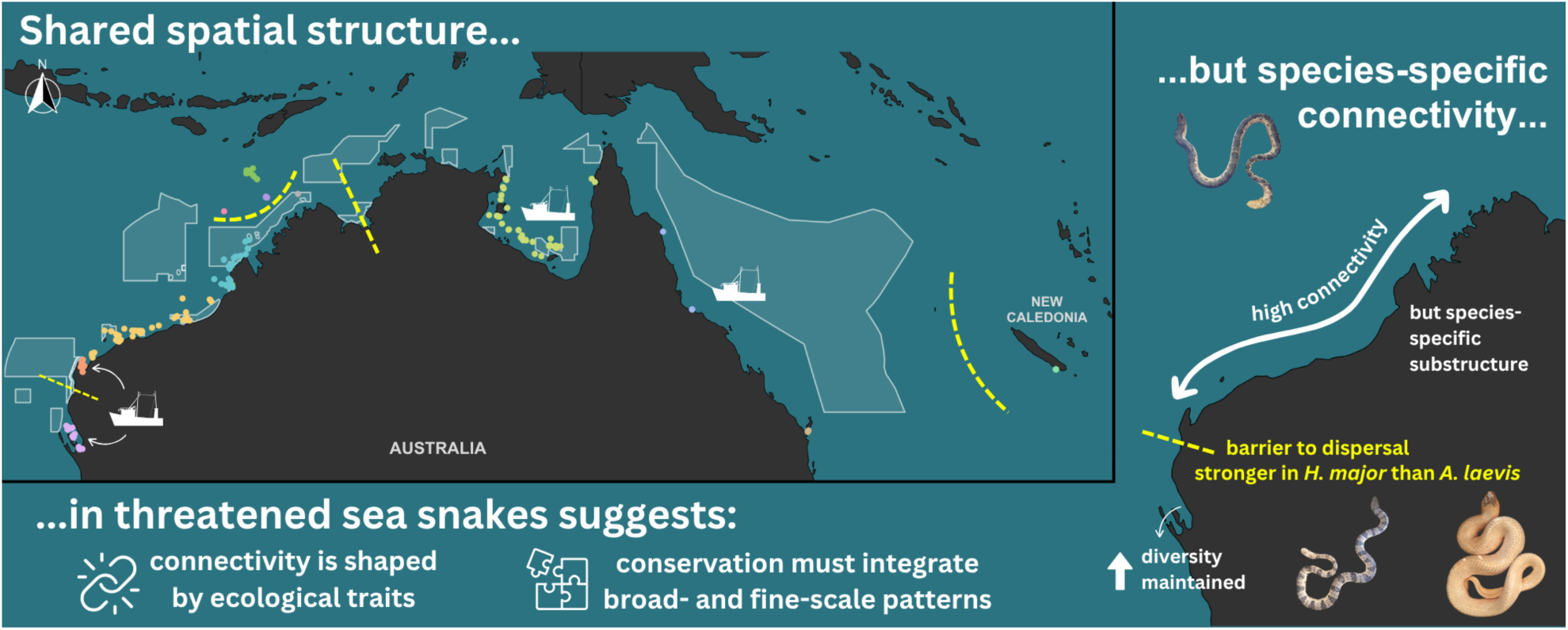

## 1. Introduction

Many marine species have expansive ranges that span marine protected areas (MPAs), commercial fishing zones, and coastal habitats threatened by infrastructure developments. Understanding spatial genetic patterns in these species is essential for their effective conservation (Dudgeon et al., 2012; Selkoe et al., 2016). In particular, analyses of inter-population connectivity can be used to determine the efficacy of MPA networks in maintaining genetic diversity and resilience, while also informing the scale and severity of threatening processes (Munguía-Vega et al., 2015). For structured populations, evaluating intrapopulation genetic diversity is crucial for setting conservation priorities. Populations with low genetic diversity may require immediate management to prevent further declines, while highly distinct populations may harbour adaptive variation critical for long-term persistence under future environmental change (Allendorf et al., 2012).

Ideally, risk assessments and recovery plans should be tailored to the needs of individual populations and species. However, while conservation genetic analyses are increasingly powerful and cost-efficient, the application of these tools is often limited for rare and elusive taxa that require logistically demanding sampling (but see Chambers et al., 2023). In some cases, well-studied species can serve as proxies for their less-understood close relatives, based on the expectation that co-occurring species with shared geographic ranges have experienced similar climatic and geological histories and should exhibit comparable spatial genetic structures (Rosauer et al., 2016; McGaughran et al., 2022). However, interspecific differences in life history, dispersal traits and ecology can also be expected to shape fine-scale phylogeographic patterns in ways that are vitally relevant for conservation (Papadopoulou & Knowles, 2016).

In this study, we examine the seascape genetic patterns of threatened Australian true sea snakes (Elapidae: Hydrophiinae)—the only fully marine reptiles alive today. These viviparous (or “true”) sea snakes evolved from terrestrial Australian elapids approximately 15 million years ago and have since developed a suite of adaptations for aquatic life, including specialised respiratory structures (Palci et al., 2019; Heatwole & Seymour, 1978), novel sensory innovations (Crowe-Riddell et al., 2016, 2019; Rossetto et al., 2023, 2024), and hydrodynamic modifications of body (Voris, 1975) and tail (Sanders et al., 2012) morphology. Sea snakes are widely distributed across shallow marine environments in the Indo-West Pacific, but northern Australia is arguably the global stronghold for these unique reptiles, supporting up to 33 species that represent all major evolutionary lineages and ecological specialisations (Rasmussen et al., 2014). In addition to their biodiversity value, sea snakes hold cultural significance for many First Nations peoples across northern Australia (McNiven, 2004).

Local declines and extirpations of sea snakes have been reported in Australia (Nowicki et al., 2019, D’Anastasi et al., 2016; Lukoschek et al., 2007) and New Caledonia (Goiran & Shine, 2013). In Australia’s Timor Sea reefs, unexplained local extinctions have led to the listing of two species as Critically Endangered (CR) on Australia’s Environment Protection and Biodiversity Conservation (EPBC) Act (1999), and one species as Endangered (EN) on both the IUCN Red List (Lukoschek et al., 2010) and EPBC Act (1999). All but one species of true sea snakes rely on coastal habitats, which often overlap with human activities including commercial fishing, industrial developments and pollution (Heatwole, 1999). A key threat to sea snakes is their incidental bycatch in Australia’s prawn trawl fisheries, where significant mortality has been documented, including for the CR-species (Kangas et al., 2015; Wassenberg et al., 1994). While the extensive commonwealth and state MPAs along Australia’s coastlines likely support sea snake conservation, the effectiveness of these protected areas will depend on their spatial alignment with the distribution and connectivity of sea snake populations.

Patterns of seascape connectivity influence both the severity of threats to sea snakes and the role of MPAs in their protection. All sea snakes give birth to live young and are relatively slow swimmers (Crowe-Riddell et al., 2024). Some species are reported to have narrow distributions, small home ranges and show high levels of philopatry (Heatwole 1999; Lukoschek et al., 2007; Lukoschek & Shine 2012; Udyawer et al., 2020). However, other species have very large geographic ranges, undergo long range movements, and occupy diverse habitat types and depths (Rasmussen et al., 2011; Udyawer et al., 2023; Sikdar et al., 2025), suggesting the potential for high population connectivity. Previous phylogeographic studies have reported varying degrees of genetic structure among species (Lukoschek et al., 2007; Nitschke et al., 2018; Garcia et al., 2022), but these studies have primarily relied on mitochondrial markers or limited nuclear data, with sparse geographic sampling for most species.

Here, we use high-density Single Nucleotide Polymorphism (SNP) analyses to investigate the population genetic diversity, connectivity and demographic history of three fully marine sea snakes (Hydrophiinae) in northern Australia and the southwest Pacific. The study species (*Hydrophis stokesii, Hydrophis major* and *Aipysursus laevis*) have similar distributions across northern Australia (Sanders et al., 2018; Crowe-Riddell et al., 2021) and are listed as Least Concern on the IUCN Red List (Guinea et al., 2010; Sanders et al., 2018; Crowe-Riddell et al., 2021). These co-occurring species are among the most commonly encountered sea snakes during surveys of rocky and soft-sediment marine habitats between 10 and 20 metre depths (Coppersmith and Sanders, pers. obs.), yet exhibit distinct morphological, ecological and life-history traits that are likely to influence patterns of connectivity and coexistence. In the case of *H. stokesii*, individuals are heavy-bodied with large heads and strong jaws, and size dimorphism is biased towards females, which can reach 2 m in length (Heatwole, 1999). This species feeds on various spiny toxic fishes (Fry et al., 2001; Wassenberg, 2001) and is distributed throughout northern Australia to south and southeast Asia (Sanders et al., 2018). Individuals of *H. major* are of moderate length and girth, with males and females reaching lengths of 1.5 m (Cogger, 2018), and this species specialises on toxic plotosid catfish across its range, from Shark Bay to New Caledonia (Shine et al., 2019; Udyawer et al., 2020). Also occurring across this range, *A. laevis* has a more generalist diet and occupies diverse habitats up to at least 100 m depths. Females of this species have very large girths and reach 1.7 m in length (Sherratt et al., 2018; Shine et al., 2023). Within *Aipysurus laevis,* the morphologically distinct Shark Bay population in Western Australia is recognised as a subspecies, *A. laevis pooleorum* (Smith, 1974). To identify common patterns and species-specific differences that are relevant to conservation managers, our sampling for these species prioritised MPAs and the major coastal trawl fishing zones.

## 2. Methods

### 2.1. Sampling, DNA extraction and sequencing

Tissue samples of *A. laevis* (n = 202), *H. major* (n = 87), and *H. stokesii* (n = 101) were collected between 1996 and 2024 from across their Australian ranges and east to New Caledonia (Table S1 and Fig. S1). Of these, 355 individuals were sampled non-destructively during fieldwork by the authors and their collaborators, and 41 were gathered from the museum collections of the Museum and Art Gallery of the Northern Territory (MAGNT), South Australian Museum (SAM) and Western Australian Museum (WAM). Tail clippings were taken from individuals collected during fieldwork by the authors and preserved in 98% ethanol. All sampling procedures were conducted in accordance with scientific licence conditions and approved animal ethics procedures.

Total genomic DNA was extracted from tail tissue, liver, and muscle samples using an adaption of the Gentra Puregene salt extraction procedure (QIAGEN, 2011) and quantified using a Quantus fluorometer. Samples with sufficient quality and quantity were sent to Diversity Arrays Technology Pty Ltd (DArT), where a proprietary genome complexity reduction pipeline was used to prepare reduced representation libraries for next generation sequencing. DNA samples were digested with the restriction enzymes *Sbfl* and *Mspl* (Kilian et al., 2012; Georges et al., 2018). The final libraries were sequenced on an Illumina Hiseq 2500 across four sequencing runs.

### 2.2. Data processing

Raw demultiplexed reads from DArT were filtered for barcode and adapter sequences using BBduk (Bushnell 2014), then screened for microbial and human contaminants using Kraken 2 (Wood et al., 2019). Unassigned reads were further filtered by Fastp (Chen, 2023) based on average quality and length, with quality metrics summarised by MultiQC (Ewels, 2016). Cleaned data were processed using ipyrad (v0.9; Eaton & Overcast, 2020) on the University of Adelaide’s Phoenix High Performance Computing (HPC) service. Sequence data for each species were aligned to their respective reference genomes (Table S2), except for *H. stokesii*, which was aligned to the *H. major* genome due to the lack of its own assembled genome. ipyrad was run with default parameters except for filter_adapters=2 and trim_reads=5,69,0,0, which apply stringent adapter and primer filtering and limit reads to a maximum of 69 base-pairs after trimming the first five bases. These settings ensured the removal of cut-site overhangs and uniform read lengths across libraries of varying lengths. Variant Call Format (VCF) files were generated for each species and were filtered on minor allele frequency (maf = 0.01) and linkage disequilibrium (ld.threshold = 0.3) using snpgdsLDpruning from the R package SNPRelate (v1.44.0) (Zheng et al., 2012). Sequencing batch effects were observed in the *H. stokesii* dataset from preliminary principal component analyses (PCA), and were managed by removing SNPs that loaded heavily on the “batch” principal component (Fig. S2). Cleaned VCF files were then converted to HDF5 format using the ipyrad Analysis Toolkit (Eaton & Overcast, 2020), specifying a custom linkage distance of 50kbp (ld_block_size=50,000) to further mitigate the effects of linkage.

### 2.3. Genetic structure and connectivity

Population genetic structure was inferred for each species using Principal Component Analysis (PCA) via the R package SNPRelate (v1.44.0) and Bayesian STRUCTURE (v2.3.4), implemented via the ipyrad-analysis toolkit. For PCA, samples within each species were grouped into populations based on collection locality (Fig. 1b, Table S3). SNPs were retained if present in at least 90% of samples included in each analysis (missing.rate = 0.1). PCA was first run with all samples to obtain broad-scale structure profiles for each species, and then separately run on samples from the Western Australian (WA) coast to investigate regional-scale population structure. Broad-scale and coastal WA analyses were filtered independently, such that SNP retention thresholds were applied separately to each dataset.

STRUCTURE analyses were used as a complementary approach to explore both broad-scale structure across all sampled populations and regional population structure within coastal WA populations. The same population groupings as the PCA were used. A burn-in of 1,000,000 iterations of the Markov Chain Monte Carlo (MCMC) were performed for each species (burnin=1,000,000), followed by 100,000 MCMC repetitions from which results were sampled (numreps=100,000). For each species, analyses were performed for *K*-values ranging from 2 to 10 for broad-scale analyses (all populations) and from 2 to 5 for regional-scale analyses (coastal WA populations), with ten independent replicates run for each value of *K* (numreps=10). The ten replicates per *K* were summarised using CLUMPP, and the most likely *K* was identified based on posterior probability and delta*K*. Q-matrices produced by STRUCTURE were exported and visualised in R (v4.1.2; R Core Team, 2021).

Differentiation between inferred population groups was quantified using Weir and Cockerham’s (1984) *F*_ST_, calculated using the snpR function *calc_basic_snp_stats* (Hemstrom & Jones, 2022). Values near zero indicate little differentiation among populations.

### 2.4. Genetic diversity

Within- and between-population diversity statistics were calculated for each species using the R package snpR (Hemstrom & Jones, 2022). Population groupings were informed by the PCA and STRUCTURE groupings above; however, diversity statistics were only calculated for populations with six or more samples, which is the minimum sample size recommended for SNP estimates of population diversity (Li et al., 2020). The snpR function *calc_basic_snp_stats* was used to calculate expected (*H*_e_) and observed (*H*_o_) heterozygosity and the inbreeding coefficient (*F*_IS_) for each population. *F*_IS_ is an index from -1 to 1 that describes genetic variation within an individual relative to its subpopulation. Values near zero meet a neutral expectation of random mating under Hardy-Weinberg expectations (Holsinger & Weir, 2009).

### 2.5. Isolation by distance across Western Australia’s continental shelf

To obtain indirect estimates of geographic population connectivity across Western Australia’s continental shelf, isolation by distance (IBD) analyses were carried out using the *mantel* function from the vegan R package (Oksanen et al., 2024). As input, pairwise Nei’s genetic distance (Nei, 1978) matrices were calculated for each species using the snpR function *calc_genetic_distances*, while pairwise least-cost (across water) distance matrices were generated from latitude and longitude data for each species using *lc.dist* function from the marmap R package (Pante & Simon-Bouhet, 2013). Spatial auto-correlation analyses were then performed using the vegan function *mantel.correlog* with 10,000 permutations, grouping the data into 200km bins.

### 2.6. Effective population size through time

Historical effective population size was estimated for each species using public whole genome resequencing data of one *H. stokesii*, three *A. laevis* and five *H. major* specimens from coastal Western Australian populations (Rossetto et al., 2024; Table S4) using the software MSMC2 v2.1.1 (Schiffels & Wang, 2020). Samples of *H. major* and *H. stokesii* were aligned to autosomal sequences from the *H. major* reference genome (GCA_033807585.1) using BWA-mem2 (Vasimuddin et al., 2019), while *A. laevis* samples were aligned to a modified *A. laevis* reference genome (GCA_040207615.1) without Z chromosome sequences. Putative Z chromosome contigs were identified in the *A. laevis* assembly by first aligning assembled contigs to the *H. major* reference genome using minimap2 (Li 2018), before calculating alignment statistics using *rustybam stats* (Vollger & Hulselmans, 2023). Contigs were removed if more than 50% of their length aligned to the Z chromosome. Following the removal of Z chromosome contigs, mappability files were generated for each reference genome using GenMap v1.3.0 (Pockrandt et al., 2020), while input files for MSMC2 were created from the alignment files using the accessory scripts *bamCaller.py* and *generate_mutlihetsep.py.* Variants were called for each chromosome/contig separately for each sample using BCFtools mpileup v1.15.1 (-q 30 -Q 30 -C 50) and BCFtools call (-c), which were processed by *bamCaller.py* to generate per-chromosome VCF and mask files. These files were then passed to *generate_multihetsep.py*, in addition to the genome mappability files, to generate the required per-sample inputs for MSMC2. MSMC2 was then run twice per-sample, first using the default clock (1*2+25*1+1*2+1*3) and once using a custom clock (2*2+25*1+2*3), with each run performing 100 EM-iterations (-i 100). MSMC2 output was then processed in R, with the results being scaled using a generation time of 10 years (Ludington & Sanders, 2020) and a mutation rate of 4.71 x 10^-9^ site/generation (Peng et al., 2020).

## 3. Results

### 3.1. Sampling, DNA extraction and sequencing

Sequencing summary statistics are presented in Supplementary Table S5. Average DArT library sizes were 1.53 million (M) (SD=256,949) for *Aipysurus laevis* samples, 1.49M (SD=345,050) for *Hydrophis major* samples and 1.75M (SD=384,047) for *Hydrophis stokesii* samples (Fig. 1a). Quality filtering resulted in 11–14.4% of reads being removed due to low quality, bacterial contamination or insufficient read-length. Reference guided ipyrad assemblies of the filtered reads produced on average 22,856 (SD=6,486) reference loci in *A. laevis* samples, 21,273 (SD=8,676) in *H. major* and 28,520 (SD=8,971) loci in *H. stokesii*, respectively housing 133,969, 52,666 and 90,080 raw genetic variants before filtering. The total number of SNPs used for each analysis is summarised in Supplementary Table S6.

### 3.2. Broad-scale genetic structure and connectivity

Analyses of broadscale population structure (PCA plots and STRUCTURE analyses including all sampled individuals) recovered several geographic groupings that were shared across species (Fig. 1; geographic groupings summarised in Fig. 1b). The PCA plots show clear differentiation between coastal populations from Western and eastern Australia, either side of the northern Kimberley, in all three species (Fig. 1c, e, g). In *H. major*, individuals from Shark Bay were differentiated from other coastal Western Australian populations on PC2. New Caledonian populations formed distinct genetic clusters in *A. laevis* and *H. majo*r (*H. stokesii* is not found in New Caledonia) but showed strongest differentiation from Australian conspecifics in *H. majo*r. Populations of *A. laevis* and *H. stokesii* from the remote Timor Sea reefs (which exclude *H. majo*r) were strongly differentiated from all other populations, including those on the Western Australian coast. One *A. laevis* was sampled from Scott Reef in the Timor Sea, and this was placed at the edge of a large cluster of Ashmore samples. There were only two samples of *H. stokesii* from the Timor Sea (both from Ashmore Reef); one of these was well separated from conspecifics on PC2, and the other was placed within the Heywood Shoal cluster. Populations of *A. laevis* and *H. stokesii* were also sampled from Heywood Shoal, which is situated on the edge of the continental shelf off the Kimberley coast of Western Australia (Fig. 1b). In both species these individuals were placed in intermediate positions between the coastal Western Australia and the offshore Timor Sea reefs, but substantially closer to the coastal populations. In all species, eastern populations were primarily sampled from the Gulf of Carpentaria (GoC). Four *A. laevis* and one *H. stokesii* sampled from North Queensland (Townsville and Cooktown, respectively) showed differentiation from Gulf of Carpentaria clusters, with *A. laevis* exhibiting subtle differentiation on PC2 and 3, whereas the *H. stokesii* individual was clearly separated from other populations along PC3 (Fig. 1c, g and Fig. S3a, c). Four *H. majo*r from a more southern locality on the south-eastern coast of Queensland (Fraser Island) also showed weak differentiation from Gulf of Carpentaria on PC1 and 3 (Fig. 1e and Fig. S3c).

Bayesian clustering analysis found the most supported number of clusters (*K*) based on the Δ*K* metric (Fig. S4) as *K* = 3 in *A. laevis* and *H. major,* and *K* = 4 in *H. stokesii*. However, inspection of ancestry profiles indicated that *K* = 4 in *H. stokesii* did not reveal an additional biologically meaningful partition, whereas *K* = 3 was consistent with observed geographic patterns. The STRUCTURE ancestry profiles were largely congruent with the PCA groupings (Fig. 1d, f, h). In *A. laevis*, four distinct geographic clusters are visible: coastal Western Australia, Timor Sea, eastern Australia (including Gulf of Carpentaria), and New Caledonia (Fig. 1d). A similar pattern was observed in *H. major*, with individuals grouping into the same three Australian regions, and Shark Bay forming a distinct cluster separate from the rest of coastal Western Australia (Fig. 1f). Individuals from south Queensland were slightly more genetically similar to those from New Caledonia, and a weak longitudinal effect was visible in the overall ancestry profile. In contrast, *H. stokesii* showed weaker population structure, with three broad clusters corresponding to coastal Western Australian, Timor Sea, and eastern Australia (Fig. 1h).

**Figure 1:**
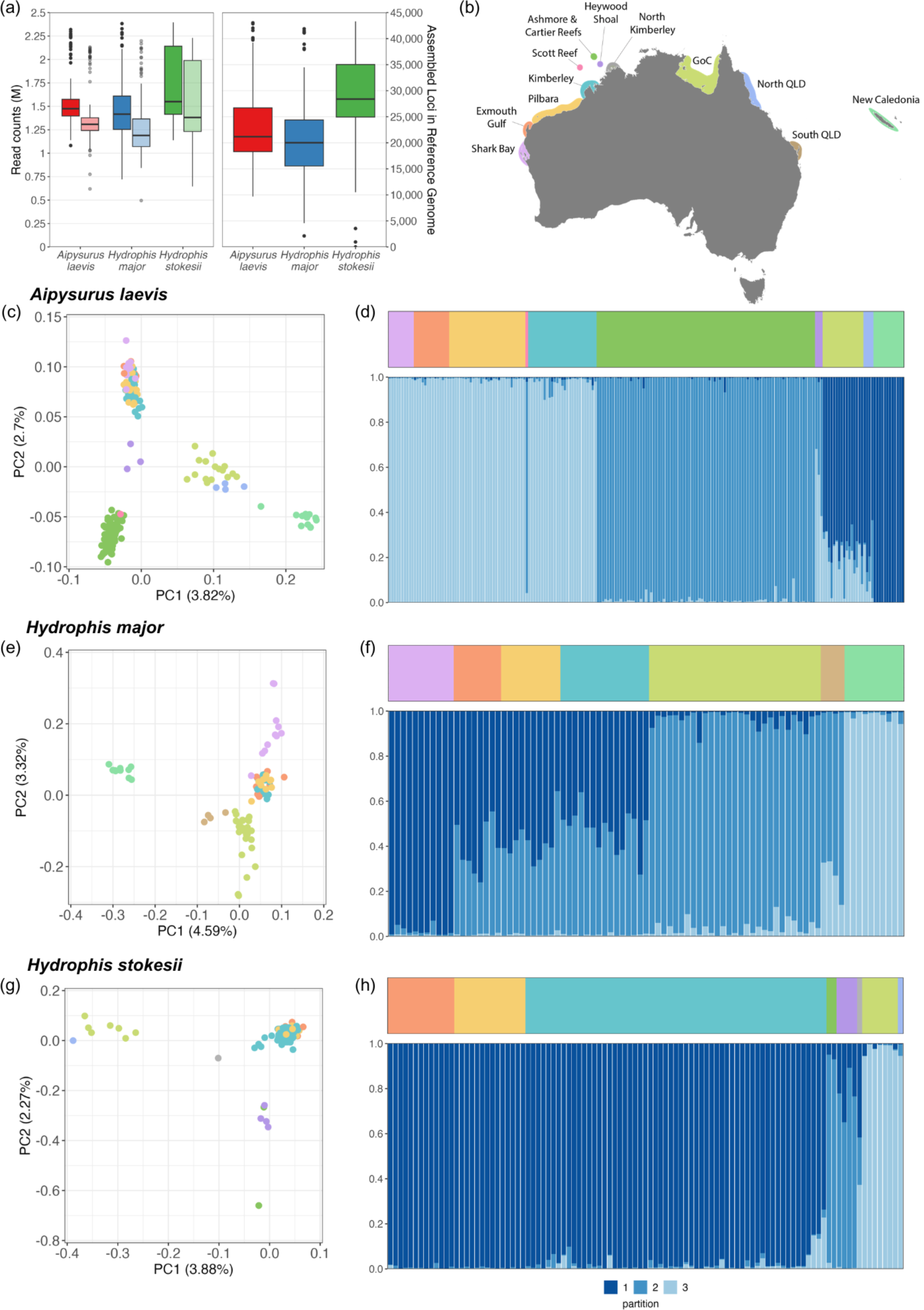
Broad-scale analyses of population genetic structure and connectivity in the three sea snake species. (a) Boxplots of read counts (left panel; raw read counts in darker shades, reads after quality control in lighter shades) and assembled loci in the reference genome (right panel). (b) Sampled regions in Australia and New Caledonia. (c, e, g) Results of Principal Components Analyses (PCA) performed on the genetic data from each species with points coloured according to the region of their origin. The values indicated along each axis correspond to the percentage of inertia explained by the corresponding axis. (d, f, h) Ancestry profiles from STRUCTURE analyses, with region of origin indicated by colour in the top panel. Regions for both STRUCTURE and PCA plots are indicated in the legend map (b).

Pairwise *F*_ST_ values and Nei’s genetic distance indicated varying degrees of genetic differentiation across populations and species (Fig. 2a, b, c). Relatively low genetic structure was observed within Western Australian populations of *A. laevis* (*F*_ST_ < 0.05 between Coastal WA, Offshore WA, and Shark Bay), suggesting ongoing gene flow. Differentiation was slightly higher between Gulf of Carpentaria and Shark Bay (*F*_ST_ = 0.11). Similarly, differentiation among Australian populations of *H. major* was low (*F*_ST_ < 0.05), except between Gulf of Carpentaria and Shark Bay (*F*_ST_ = 0.099). Australian sub-populations in both species, however, were strongly differentiated from New Caledonian populations (*F*_ST_ = 0.16–0.25), which also exhibited lower genetic diversities (Table 1). Higher differentiation was detected in *H. major* between Shark Bay and Coastal WA (0.048) compared to *A. laevis* (0.011). In contrast, *H. stokesii* showed high differentiation between the Gulf of Carpentaria and both the Offshore WA (0.22) and Coastal WA (0.14) populations.

Nei’s genetic distance followed a similar pattern, with the greatest distances observed between New Caledonia and all other populations in *A. laevis* and *H. major*, and between Gulf of Carpentaria and all other populations in *H. stokesii*. Populations with low *F*_ST_ values, such as those within Western Australia, exhibited lower Nei’s genetic distances, further supporting high levels of genetic connectivity in these ranges. These results suggest that geographic barriers and population isolation contribute to genetic structuring, particularly in New Caledonia and Gulf of Carpentaria.

**Figure 2:**
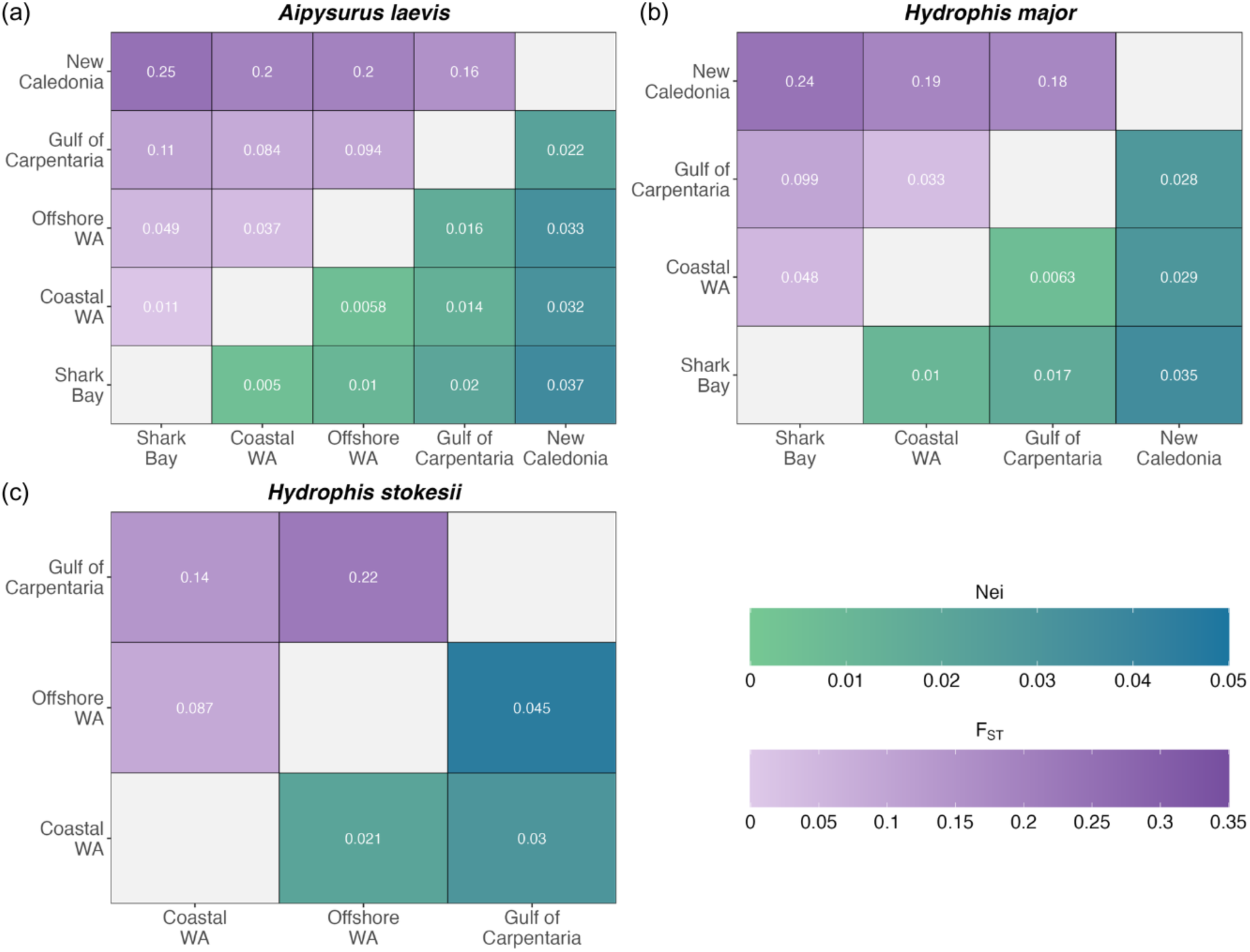
Pairwise Nei’s genetic distance (shades of green), and *F*_ST_ (shades of purple) comparisons among population groupings with 6 or more individuals for (a) *A. laevis*, (b) *H. major*, and (c) *H. stokesii*.

### 3.3. Regional genetic structure

Fine-scale population structure was analysed separately for the Western Australian coastal localities, which were mostly undifferentiated in the broadscale analyses. PCA and STRUCTURE analyses further differentiated geographic patterns within these coastal localities (Fig. 3A; S5). In the PCAs, the strongest differentiation (along PC1) was seen between populations of *A. laevis* and *H. majo*r from Shark Bay and the remainder of the Western Australian coast (*H. stokesii* is not found in Shark Bay) (Fig. 3A-i,ii). PC2 in *A. laevis* shows a mostly undifferentiated cluster of samples from the Exmouth Gulf, Pilbara and Kimberley. PC2 in *H. majo*r shows a similarly undifferentiated cluster from the Exmouth Gulf, Pilbara and Kimberley, with the Kimberley samples having the greatest spread along this axis. Finally, PC1 in *H. stokesii* shows a slight longitudinal effect, with the greatest differentiation seen in the North Kimberly sample and other coastal populations (Fig. 3A-iii).

Inspection of the STRUCTURE ancestry profiles revealed that at *K* = 3 (the optimal number of clusters for all three species; Fig. S4) revealed weak fine-scale structure among Western Australian coastal populations (Fig. S5). In *A. laevis*, Shark Bay individuals showed subtle shifts in ancestry proportions (∼10%), indicating limited differentiation. In contrast, *H. major* exhibited clearer structure, with Shark Bay individuals showing a greater contribution from a secondary cluster (∼5–30%) compared to northern populations. No structure was apparent in *H. stokesii* across any value of *K*.

Mantel tests excluded Shark Bay samples due to their clear genetic differentiation from northern coastal populations. Results indicated a weakly positive and significant IBD relationship between genetic distance (Nei) and geographical least-cost distance (km) for *A. laevis* (*r*^2^ = 0.06, *p* = 0.046; Fig. 3B-i). In contrast, *H. major* and *H. stokesii* showed no significant IBD. Spatial autocorrelation analyses revealed mostly neutral Mantel autocorrelations across distance classes in both *A. laevis* and *H. major*, indicating little evidence of spatial genetic structure among these coastal localities (Fig. 3C-i,ii). In *H. stokesii*, a positive correlation was observed between individuals separated by <500 km, with negative correlations at larger distances (∼1,500 km), while other distance classes showed values close to zero (Fig. 3C-iii).

**Figure 3:**
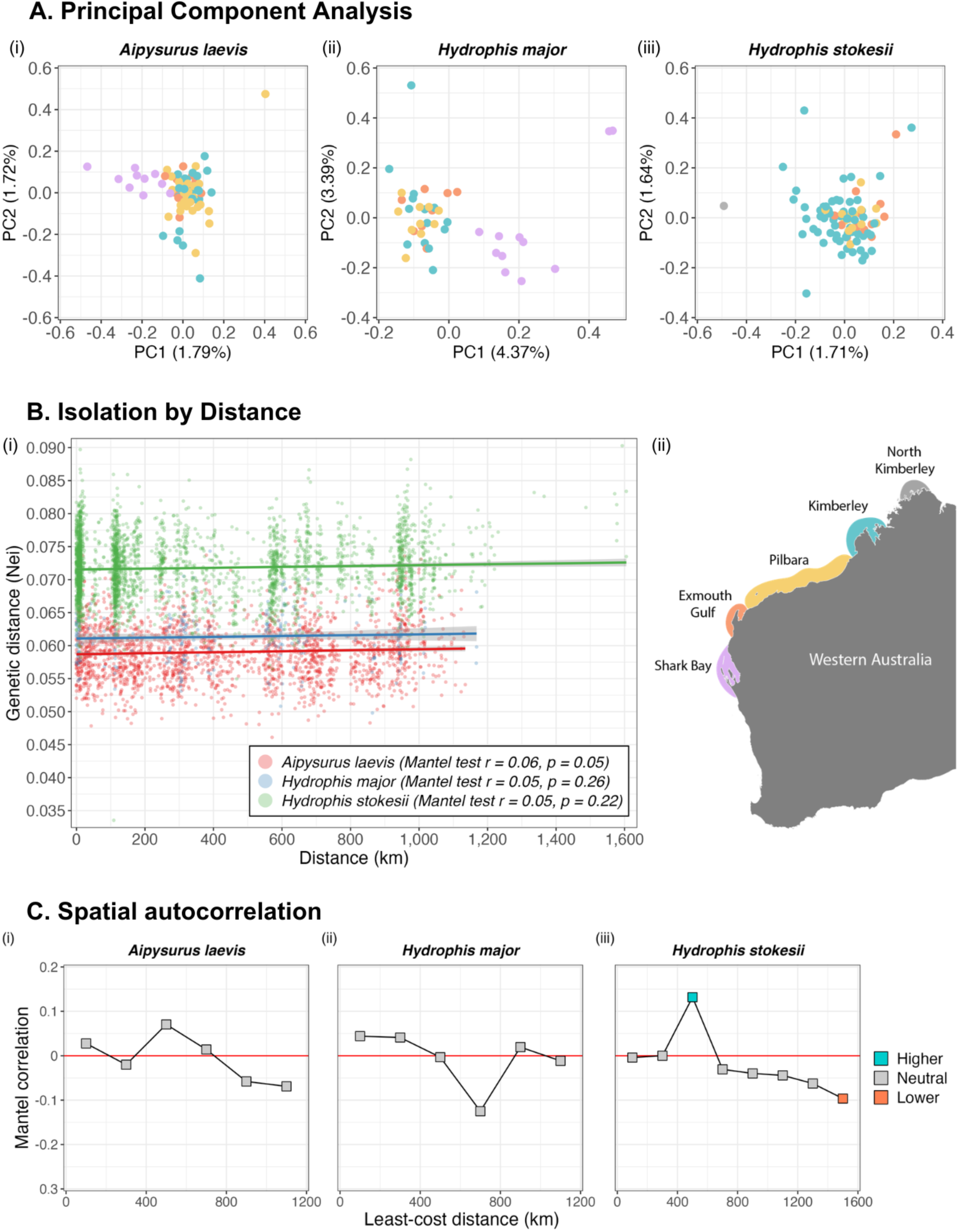
**A**. Results of Principal Components Analyses (PCA) performed on the genetic data from i) *A. laevis*, ii) *H. major*, and iii) *H. stokesii* sampled across their coastal Western Australian ranges. The values indicated along each axis correspond to the percentage of inertia explained by the corresponding axis. **B.**i) Isolation by distance plot from Mantel tests comparing population pairwise Nei’s genetic distance and geographical distance (km) for each sea snake species (excluding Shark Bay samples due to their genetic differentiation from other coastal populations), and ii) Map showing coastal WA population groupings in which the PCA points in **A** are coloured. **C.** Spatial autocorrelation correlograms for i) *A. laevis*, ii) *H. major*, and iii) *H. stokesii* comparing Mantel correlations between individual pairwise Nei’s genetic distance at increasing distance classes (km).

### 3.4. Genetic diversity

Genetic diversity varied among species and across the genetic populations identified in the PCAs (Table 1). Nucleotide diversity (*π*) was highest in *H. stokesii*, ranging from 0.101 in Gulf of Carpentaria to 0.127 in Coastal WA. *A. laevis* exhibited slightly lower nucleotide diversity (0.072– 0.115), while *H. major* had the lowest values overall (0.071–0.109). In both *A. laevis* and *H. major*, New Caledonian populations exhibited the lowest nucleotide diversity for each species (*H. stokesii* is not found in New Caledonia).

Observed heterozygosity (*H*_o_) broadly matched expected heterozygosity (*H*_e_) in most populations, although *H*_o_ was generally slightly lower. Populations of *H. stokesii* exhibited the highest heterozygosity of the three species (*H*_o_ = 0.090–0.121), followed by *A. laevis* (0.068–0.110) and *H. major* (0.069–0.104). These patterns were broadly consistent with trends in nucleotide diversity.

Inbreeding coefficients (*F*_IS_) varied across populations and species. Most *A. laevis* populations showed low *F*_IS_ values (0.024–0.056), with the exception of the Gulf of Carpentaria where values were markedly higher (0.319), consistent with local substructure or non-random mating. Most *H. major* populations showed similarly low *F*_IS_ values (0.033–0.046), with a slight elevation in the Gulf of Carpentaria (0.073). In contrast, *H. stokesii* exhibited higher *F*_IS_ values overall, particularly in Offshore WA (0.117) and the Gulf of Carpentaria (0.123). In both *A. laevis* and *H. major*, *F*_IS_ values were lowest in New Caledonia.

**Table 1:**
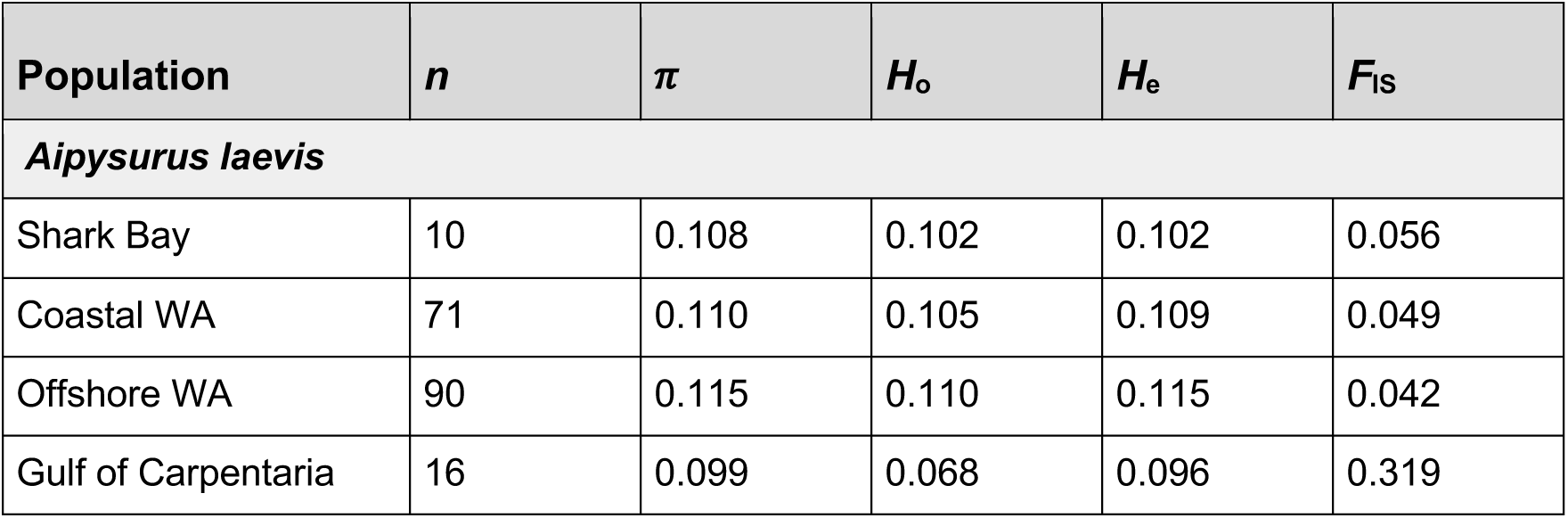
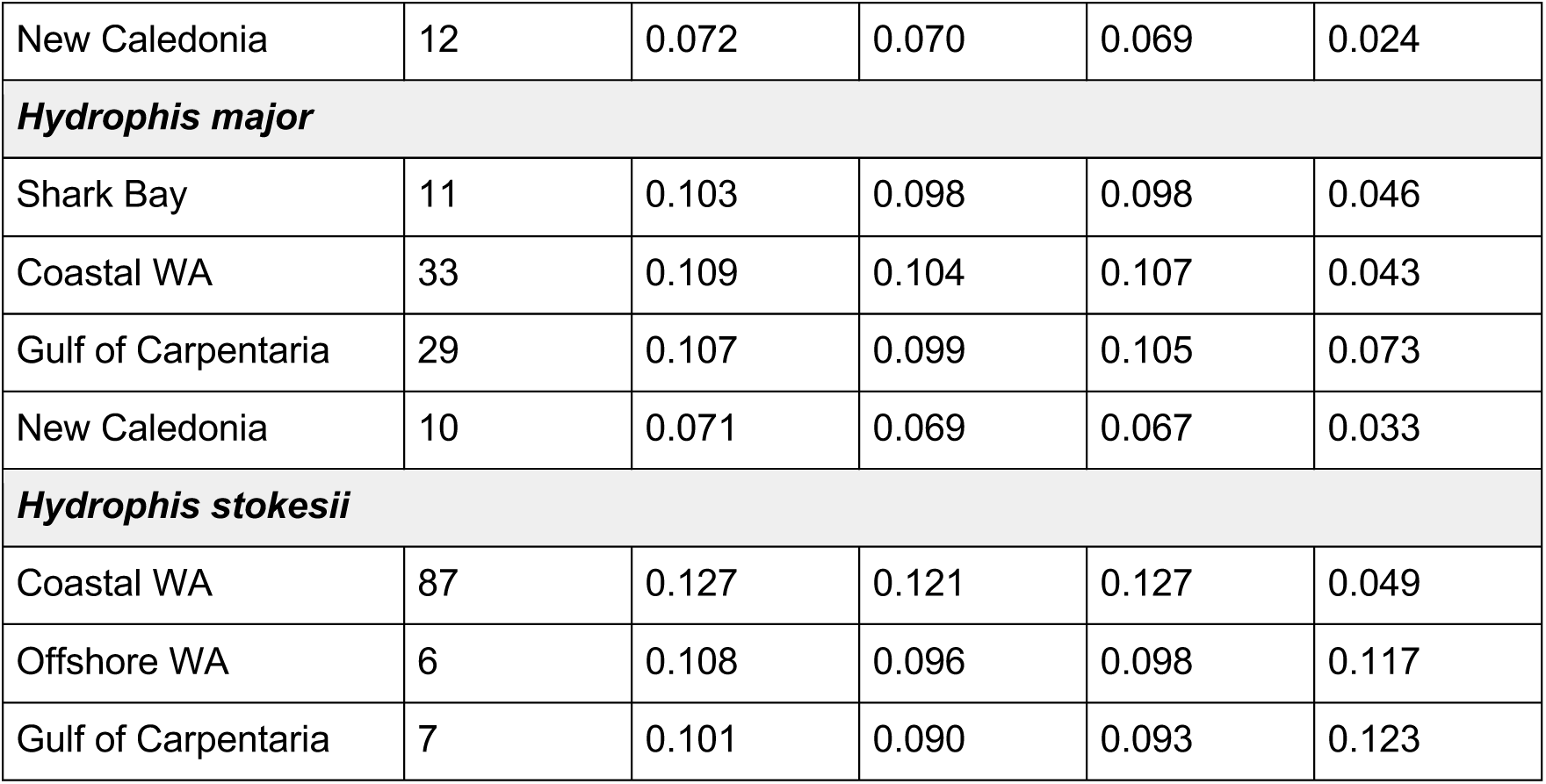
Genetic diversity estimates and sample sizes for each genetic cluster for *Aipysurus laevis*, *Hydrophis major*, and *Hydrophis stokesii*. *n* = sample size, *π* = nucleotide diversity (pixy), *H*_o_ = observed heterozygosity, *H*_e_ = expected heterozygosity, *F*_IS_ = inbreeding coefficient. Metadata associated with each site is shown in Table S1.

| Population | $n$ | $\pi$ | $H_o$ | $H_e$ | $F_{IS}$ |
| --- | --- | --- | --- | --- | --- |
| ***Aipysurus laevis*** |  |  |  |  |  |
| Shark Bay | 10 | 0.108 | 0.102 | 0.102 | 0.056 |
| Coastal WA | 71 | 0.110 | 0.105 | 0.109 | 0.049 |
| Offshore WA | 90 | 0.115 | 0.110 | 0.115 | 0.042 |
| Gulf of Carpentaria | 16 | 0.099 | 0.068 | 0.096 | 0.319 |
| New Caledonia | 12 | 0.072 | 0.070 | 0.069 | 0.024 |
| ***Hydrophis major*** |  |  |  |  |  |
| Shark Bay | 11 | 0.103 | 0.098 | 0.098 | 0.046 |
| Coastal WA | 33 | 0.109 | 0.104 | 0.107 | 0.043 |
| Gulf of Carpentaria | 29 | 0.107 | 0.099 | 0.105 | 0.073 |
| New Caledonia | 10 | 0.071 | 0.069 | 0.067 | 0.033 |
| ***Hydrophis stokesii*** |  |  |  |  |  |
| Coastal WA | 87 | 0.127 | 0.121 | 0.127 | 0.049 |
| Offshore WA | 6 | 0.108 | 0.096 | 0.098 | 0.117 |
| Gulf of Carpentaria | 7 | 0.101 | 0.090 | 0.093 | 0.123 |

### 3.5. Effective population size through time

MSMC2 analyses reconstructed effective population sizes (*N*_e_) over time segments that included the period of shared ancestry among the three species (Fig. 4). Interspecific variation in *N*_e_ trajectories was highest in the oldest and most recent time intervals, when fewest coalescent events are available for reconstructing coalescence rates. All species exhibit fluctuations in *N*_e_, but these are least prominent in the *H. stokesii* genome. All three species show lowest *N*_e_ at intermediate time segments. The three *A. laevis* and four *H. major* show clear upticks and declines in *N*_e_, and all three species show an increase to peak levels of *N*_e_ in the most recent time segment. Individual estimates of *N*_e_ in the most recent time segment overlap among the three species, but are highest in *A. laevis,* followed by *H. major,* then *H. stokesii*.

**Figure 4:**
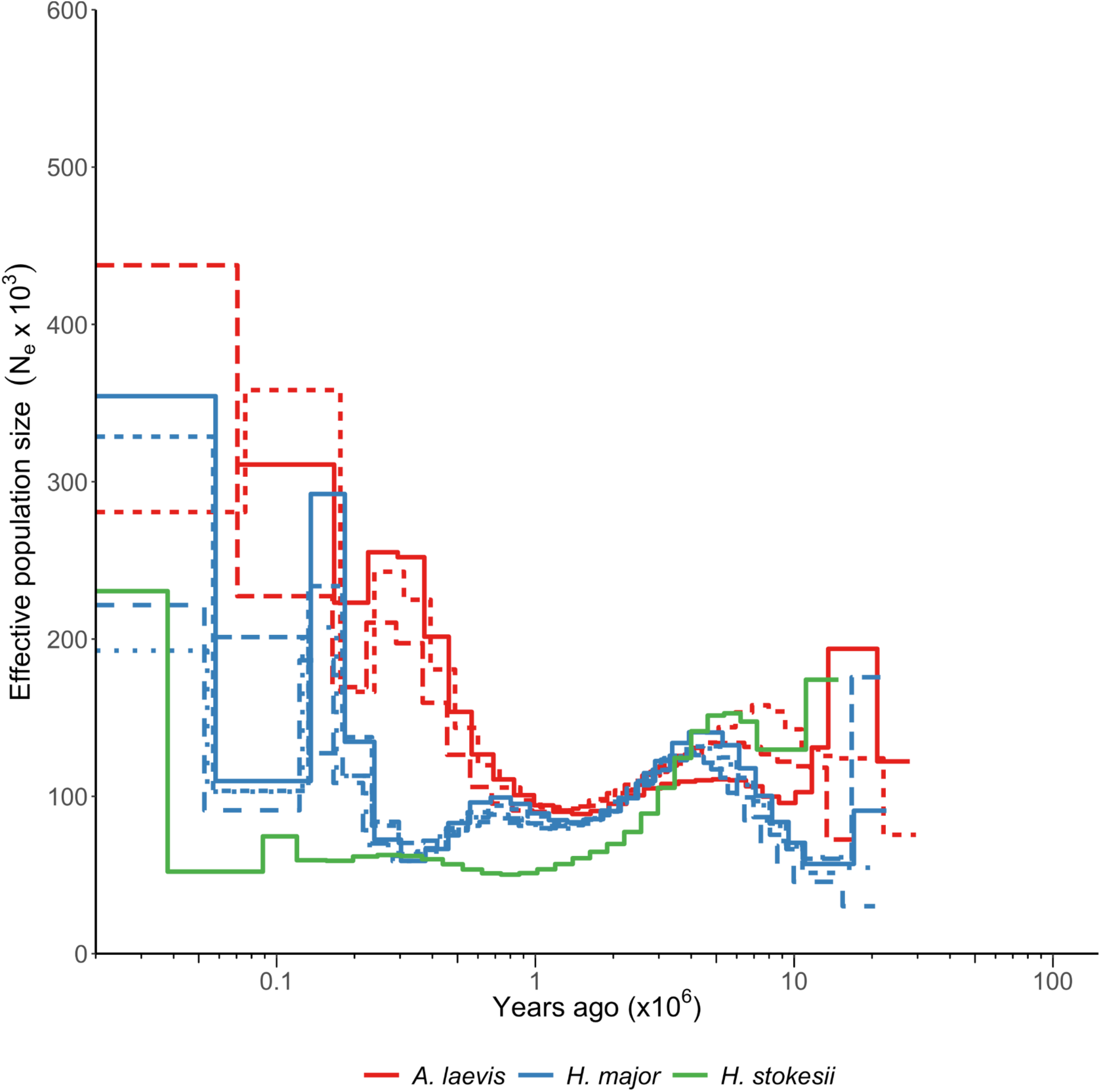
Historical effective population sizes reconstructed using MSMC2 for A. laevis, H. major and H. stokesii. Time and Ne are scaled using a generation time of 10 years and mutation rate of 4.71x10^-9^ per-site/per-generation. The molecular clock ‘2*2+25*1+2*3’ is used, with the first two interval parameters spanning two intervals, the next 25 parameters spanning one interval each, and the last two parameters each spanning 3 intervals. This clock was used to reduce aberrant Ne estimates at extremely recent and distant time points.

## 4. Discussion

This study used reduced-representation genotyping and whole genome sequences to investigate population genetic patterns in three co-occurring sea snake species across northern Australia to New Caledonia. As one of the first studies to examine the population genetics of sea snakes in this region, our findings contribute to a broader understanding of population connectivity in the remote waters of northwestern Australia and the southwest Pacific. Our analyses revealed congruent population structure among species at broad geographic scales, as well as species-specific differences in putative connectivity at finer temporal and spatial scales (Fig. 5). These results are discussed below in the contexts of recent phylogeographic history and spatial planning for sea snake conservation.

**Figure 5:**
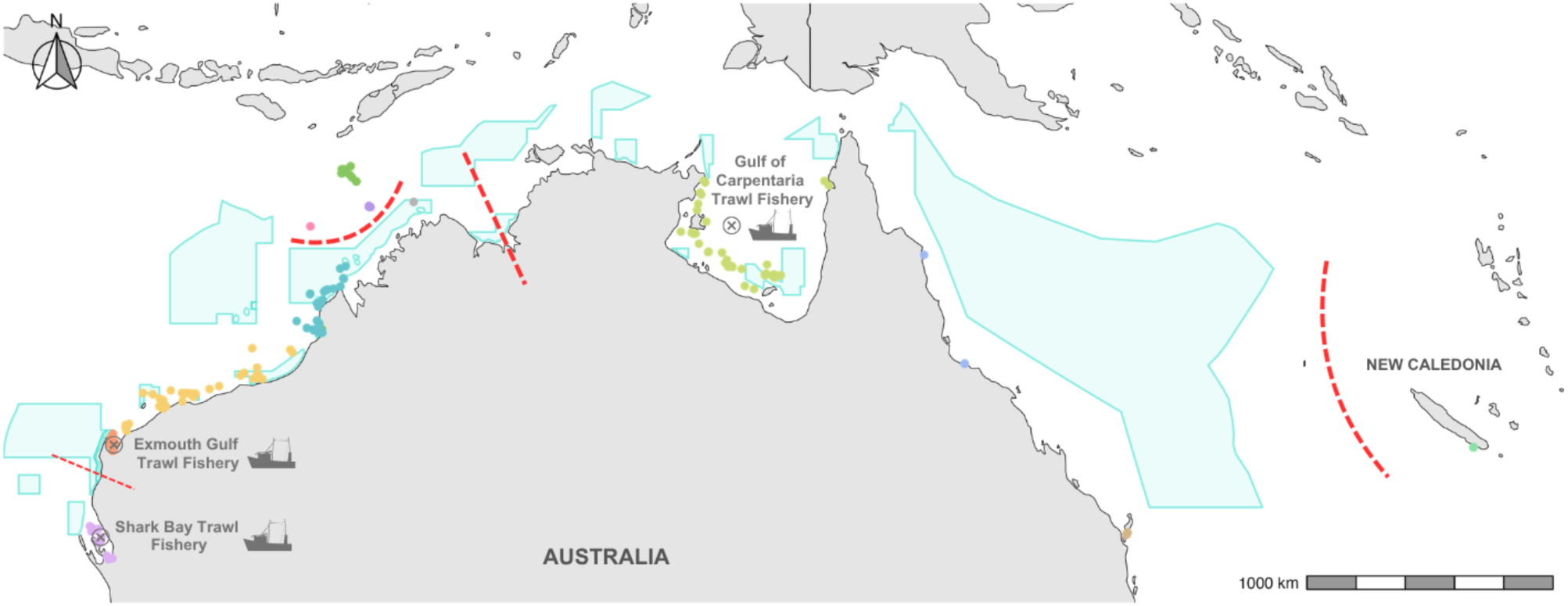
Map showing sampling for all species (coloured by populations used in our analyses), shared major genetic breaks (thick red dashed lines) and minor genetic breaks (thin red dashed line), along with Australian MPAs (shaded blue polygons), and the three major prawn fisheries in north-western Australia (boat icons).

### 4.1. Discontinuities across Australia and the southwest Pacific

Broadscale population genetic comparisons revealed common phylogeographic patterns across species, with major genetic breaks aligning with known historical biogeographic barriers to connectivity in marine taxa. The most pronounced genetic break was observed between Australian and New Caledonian populations of *A. laevis* and *H. major* (*H. stokesii* is not found in New Caledonia) (Fig. 1). In both species, individuals from New Caledonia formed discrete genetic clusters that were clearly separated from Australian populations but showed highest affinity with eastern Australian samples. This genetic split aligns with a substantial dispersal barrier across the Coral Sea, including ∼1,330 km of open-water and a deep-sea trench that must limit gene flow in sea snakes. Similar genetic breaks across the Coral Sea have been documented in numerous other marine taxa, including sharks (Giles et al., 2014), manta rays (Lassauce et al., 2022), reef fishes (Planes & Fauvelot, 2002), and dugongs (Verger et al., 2025). Interestingly, genetic differentiation was particularly pronounced in *H. major,* with the New Caledonian population of this species also exhibiting lower nucleotide diversity and heterozygosity than its Australian counterparts. The New Caledonian *A. laevis* population also exhibited the lowest genetic variation of any sampled population of this species. However, the absence of inbreeding (low *F*_IS_) signals in New Caledonian sea snakes suggests that they have maintained stable, outbreeding populations despite their reduced genetic diversity. Hence, while it is possible that New Caledonian populations experienced historical bottlenecks associated with founder events, an equally plausible explanation for their relatively low diversity is their smaller, isolated distribution in comparison to the very wide ranging and substructured Australian sea snake populations. Our Markovian coalescent analyses of genome sequences for Western Australian populations revealed population expansions in all three species within the last ∼10,000 years. This timing coincides with sea level rise during the current interglacial period, when inundation of Australia’s continental shelf reconnected shallow marine habitats. Increased habitat availability and connectivity have been linked to similar genetic signals of recent population expansion and reconnection in shallow marine fishes and invertebrates on Australia’s continental shelf (Ludt & Rocha, 2015; Payet et al., 2024).

Within Australia, we identified a pronounced split between northwestern (coastal WA and Timor Sea) and northeastern (Gulf of Carpentaria to southern Queensland) localities for all species (Fig. 1). While gaps in sampling between the Kimberley and the Gulf of Carpentaria may have contributed to this separation, the comparably high genetic connectivity observed across ∼1,300 km of the northwestern continental shelf suggests that this result is not merely an artifact of incomplete sampling. Genetic discontinuities across the Torres Strait have been documented in a wide range of marine taxa, from sessile invertebrates (Mirams et al., 2011), to large fishes (e.g., Veilleux et al., 2011; Taillebois et al., 2017, Barton et al., 2018, Loughnan et al., 2019), turtles (Jensen et al., 2019; FitzSimmons et al., 2020) and dugongs (Blair et al., 2014). These patterns are often attributed to the historical land barrier that intermittently connected Australia and New Guinea during Plio-Pleistocene sea level fluctuations (Voris, 2000). We also found notable structure between Gulf of Carpentaria and southeast Queensland populations in *H. major* (Fig. 1e, f), a pattern which has been reported in genetic assessments of other sea snake species (Garcia et al., 2022). However, our restricted sampling across Queensland may have contributed to the apparent genetic separation of clusters. We recommend further sampling to resolve population connectivity along the east coast, particularly given the Queensland East Coast Otter Trawl Fishery captures over 100,000 sea snakes as bycatch annually (Courtney et al., 2010). Northeastern (Gulf of Carpentaria) and northwestern populations of each species showed similar patterns of genetic variation, across metrics, but with consistently higher heterozygosities in Australian populations of *H. major*, and relatively high *F*_IS_ in the Gulf of Carpentaria population of *A. laevis*. *F*_IS_ was, in fact, highest in the Gulf of Carpentaria across all species, suggesting increased population substructure or non-random mating in this region. This pattern may reflect admixture among historically isolated populations following the submergence of the Torres Strait land bridge, although it is also possible that fine-scale structure is maintained within this region.

The strong genetic divergence between both northwestern and northeastern coastal populations within Australia suggests a history of isolation that persists to the present day, supporting the recognition of these regions as distinct conservation units. These results highlight the need for independent management strategies tailored to each region. However, as this genetic break overlaps with gaps in our sampling, future studies should incorporate more extensive sampling between the Kimberley and the Gulf of Carpentaria to better delineate the boundaries of these conservation units.

### 4.2. Isolation of threatened Timor Sea populations

The final genetic break identified in our broadscale analyses occurred between offshore Timor Sea populations (including Ashmore, Cartier, Hibernia and Scott reefs) and coastal WA populations (continental shelf localities between Shark Bay and the Kimberley). Clear breaks in connectivity were observed in both *A. laevis* and *H. stokesii* (*H. major* is not found in the offshore Timor Sea reefs), with minimal shared ancestry between offshore and coastal populations of this species. These convergent genetic splits likely reflect shared responses to a combination of historical barriers associated with sea level fluctuations, and contemporary oceanographic features that delimit the Canning and Kimberley bioregions (Pope et al., 2015). The magnitude of genetic divergence between coastal and offshore populations suggests long-term historical isolation that has persisted to the present day (Nankivell and Sanders, 2025).

The strong genetic differentiation observed in offshore Timor Sea reef populations highlights their conservation importance. These small, isolated populations are particularly vulnerable to environmental disturbances and have experienced unexplained population declines at Ashmore Reef over decadal timescales (D’Anastasi et al., 2016; Lukoschek et al., 2007; Sanders et al., 2015). Recent sightings of nine species (including *A. laevis*) in mesophotic habitats on the outer edge of Ashmore Reef suggest a potential recovery of sea snake populations at Ashmore (Speed et al., 2022). However, our findings indicate that recolonisation from coastal populations is unlikely, as *A. laevis* and *H. stokesii* show minimal recent shared ancestry with mainland populations. Tissue sampling for genetic analyses should be collected to determine the most likely origin of these deeper water snakes.

The relatively high genetic diversity observed in these offshore populations suggests several possibilities. It is possible that these populations are sustained by connectivity within the Timor Sea itself. Timor Sea populations (sampled before the documented declines at Ashmore Marine Park) exhibited nucleotide diversity and heterozygosity estimates comparable to those of the geographically much larger conspecific populations on the continental shelf, particularly in *A. laevis* (Table 1). This could indicate that even the most isolated sea snake populations remain genetically diverse. Alternatively, genetic diversity might be maintained through connectivity with nearby shoal systems that have yet to be surveyed for sea snakes. For *H. stokesii*, occasional migrants from South-East Asian populations may also contribute to maintaining diversity in the Timor Sea, whereas *A. laevis* does not occur further north and therefore lacks this potential source of gene flow.

Both species occurring at Heywood Shoal (*A. laevis* and *H. stokesii*), an outer shelf shoal situated between the Kimberley coast and Ashmore and Cartier Reefs, were genetically distinct from sampled conspecifics, exhibiting both mixed, and unique ancestry from both coastal and offshore populations and occupying an intermediate position in the PCAs (Fig. 1c, d, g, h). Although our limited sampling in this region precludes definitive conclusions on fine-scale connectivity between these offshore reefs, this finding suggests that the barrier to dispersal between Ashmore & Cartier Reefs and Heywood Shoal is stronger than that observed between Ashmore and the single Scott Reef sample, despite the deeper waters and greater distance that separates those reefs. Understanding the extent of gene flow among these isolated reef systems should be a priority for future research, as it may inform conservation strategies aimed at ensuring the long-term viability of these highly fragmented and unique populations. In particular, it will be crucial to determine whether sea snake populations on Ashmore and Cartier Reefs are connected to each other, and to adjacent shoal systems and whether the recent rediscovery of sea snakes on the mesophotic edges of these reefs (Speed et al., 2022) reflects a relict population, *in situ* recovery from a bottlenecked population, or recruitment from extralimital (e.g., shoal) populations. An apparent lack of recovery in the shallows at Ashmore reef raises questions over whether recovery via connectivity with adjacent populations occurs over ecologically relevant timescales.

### 4.3. Genetic connectivity of the northwestern continental shelf

Analyses at fine spatial scales revealed important patterns that were not detected in the broadscale analyses. A key finding was the extensive population connectivity observed along ∼1,300 km of the northwestern continental shelf, between Exmouth and the Kimberley. This was expected given the vast expanse of continuous shallow habitat in this region and the absence of major biogeographic barriers. In fish, high connectivity across the northwestern Australian shelf is thought to be facilitated by the strong Holloway and Leeuwin currents (D’Adamo et al., 2009), which carry larvae distances of up to 1,000 km (Coulson et al., 2023). Although sea snakes are viviparous and lack a larval phase, ocean currents must still play a role in their seascape movement during both juvenile and adult life stages, as sea snakes exhibit daily individual movements to forage and breathe. The predominantly southwesterly flow of these currents could create a directional bias in connectivity, facilitating southward dispersal along the continental shelf while potentially restricting northward gene flow. However, it is important to note that patterns of genetic connectivity do not equate to contemporary demographic connectivity, including the rate at which migrants contribute to population replenishment following local declines. Complementary studies of sea snake movement ecology using telemetry (e.g. Coppersmith et al. 2025) are needed to quantify functional connectivity in sea snakes.

Consistent with the largely undifferentiated clusters observed between Exmouth and the Kimberley, Mantel tests and spatial autocorrelation analyses showed little evidence for isolation by distance patterns. Although a weak positive correlation was found between genetic differentiation and geographic distance in *A. laevis* (Fig. 3B), no significant isolation by distance was observed in *H. major* or *H. stokesii*, and spatial autocorrelation analyses revealed mostly neutral patterns across distance classes, indicating limited spatial genetic structure at this scale. Together, these results suggest generally high connectivity along the northwestern continental shelf, with only weak signatures of isolation by distance in *A. laevis*. These subtle differences nevertheless highlight the importance of species-specific traits in spatial planning for conservation (Papadopoulou & Knowles, 2016). All three study species are large-bodied species, thus likely to have comparable swimming abilities; however, they exhibit clear ecological differences. A fish generalist, *A. laevis* occupies diverse habitats across the northwestern continental shelf, from coral reefs to unstructured deep-water environments exceeding 100 m (Speed et al., 2022). In contrast, *H. stokesii* and *H. major* are largely restricted to shallow habitats (<50 m) and specialise in spiny-toxic fish prey such as porcupine fish (Diodontidae), toadfish (Batrachoididae) and frogfish (Lophiiformes) (Voris & Voris, 1983, Fry et al., 2001), and eel-tailed catfish (Plotosidae) (Shine et al., 2019; Udyawer et al., 2020), respectively. These habitat and dietary preferences may shape population abundance, distribution, and maximum dispersal distances, influencing connectivity at fine geographic scales. Even rare long-distance dispersal events could significantly impact local connectivity (e.g. Macleod et al., 2024). Studies of sea snake movement patterns and dispersal abilities are needed to understand how species-specific differences in connectivity are shaped by behavioural, ecological and broader phylogeographic influences.

A notable exception to this regional connectivity was the genetic discontinuity between Shark Bay and northern localities, including Exmouth Gulf. This genetic break aligns with a well-documented biogeographic boundary near Point Cloates, just south of Ningaloo Reef, and is associated with distinct oceanographic features (Woo et al., 2006; Spalding et al., 2007). Shark Bay represents the southern limit of sea snake distributions and supports five resident species (Storr & Harold, 1978), including *A. laevis* and *H. major*, both of which exhibited some degree of genetic isolation. This discontinuity is likely driven by a combination of the strong, southward-flowing Leeuwin Current, the dominant boundary current of the Western Australian coastline (Godfrey & Ridgway 1985), and the limited extent of shallow habitat along the narrow Ningaloo coastline (Woo et al., 2006). Although the Leeuwin Current promotes southward connectivity along the broad northwestern shelf, hydrodynamic studies show that it shifts offshore south of Point Cloates, while a weaker, north-flowing nearshore Ningaloo Current forms (Woo et al., 2006; Xu et al., 2013). As a result, the Leeuwin Current largely bypasses the entrance of Shark Bay rather than flowing into it, limiting dispersal into the embayment. Similar genetic breaks have been observed in a range of other marine taxa, including reef fishes (DiBattista et al., 2017), corals (Thomas et al., 2017), crabs (Briggs et al., 2024), clams, and mangroves (Johnson et al., 1993).

Differentiation between Shark Bay and northern populations was weakest in *A. laevis*, with low fixation indices but a discrete cluster of Shark Bay versus Exmouth and Pilbara samples. In contrast, *H. major* showed a much higher fixation index and formed a distinct genetic cluster even in broadscale analyses that included samples from eastern Australia and New Caledonia (Fig. 1e). Notably, the *A. laevis* population in Shark Bay is recognised as a separate subspecies restricted to this region, *Aipysurus laevis poolerum* (Smith, 1974), which is distinguished from the nominotypical *Aipysurus laevis laevis* found throughout northern and eastern Australia by its generally darker colouration and smaller body size. Our finding of relatively weak differentiation in a recognised subspecies (*A. l. pooleorum*), versus cryptic genetic isolation in *H. major*, highlights the need for genetic assessments that account for both subspecies- and population-level connectivity to guide conservation priorities in sea snakes.

Despite their apparent isolation, Shark Bay populations maintained relatively high genetic diversity, potentially reflecting historical connectivity with northern populations and occasional contemporary gene flow. However, limited connectivity suggests that any population declines in Shark Bay would not be rapidly replenished from northern source populations. This has important implications for fisheries management, as sea snakes are frequently caught as bycatch in bottom-trawling operations across their range. Three major prawn fisheries operate in Western and northern Australia—in Shark Bay, Exmouth Gulf, and the Gulf of Carpentaria—all of which have reported substantial sea snake bycatch. For instance, the Shark Bay Prawn Managed Fishery and Exmouth Gulf Prawn Managed Fishery reported 3,168 and 871 interactions with sea snakes, respectively, in the 2021/2022 season (Newman et al., 2023), while the Northern Prawn Fishery (including the Gulf of Carpentaria) documented 12,643 sea snakes caught in 2023 (Meteyard, 2024). Given the implied reduction of gene flow into Shark Bay, the sea snake populations assessed here may be particularly vulnerable to declines, highlighting the need for localised conservation measures. Similarly, the genetic break between the Gulf of Carpentaria and Western Australian populations suggests that bycatch impacts should be assessed independently for each region.

### 4.4 Conclusions

Our findings reveal both shared and species-specific patterns of population genetic differentiation among three sea snake species in northern Australia, providing valuable insights for conservation management. Pronounced genetic breaks between northwestern and northeastern Australia, as well as northwestern coastal and offshore populations, indicate long-term isolation in response to shared biogeographic barriers. These findings, along with subtler genetic differentiation of Shark Bay populations, support the delineation of conservation units for sea snakes that will enable region-specific management strategies. Large, genetically diverse populations with relatively high connectivity across the northwestern continental shelf were found in all three species. This connectivity, coupled with their presence in the extensive Australian Marine Park networks, may enhance resilience to localised habitat loss and population fluctuations. Consequently, regional-scale threats such as ocean warming, extreme weather events, and large-scale habitat degradation should be considered the primary risks to sea snake populations in this region. Finally, while all three species show congruent population genetic structure in response to historical land barriers and contemporary oceanographic features, they exhibit subtle differences in connectivity across the continuous habitats of the northwestern continental shelf. We suggest that these differences are shaped by species-specific ecological specialisation and movement behaviours that influence connectivity at finer temporal and spatial scales. Taken together, our results highlight the importance of shared phylogeographic patterns as well as species-specific processes in spatial planning for sea snake conservation.

## Data availability

The processed datasets, analysis and visualisation scripts, and files required to reproduce the analyses are available on GitHub (https://github.com/a-lud/sea-snake-dart).

## Animal Ethics

All sampling procedures were in accordance with methods approved by the University of Adelaide’s Animal Ethics committee (permit 34903), James Cook University (permit A2310), Charles Darwin University Animal Ethics (permit A18029). Collection permits were granted from the Western Australian Department of Biodiversity, Conservation and Attractions (FO25000393-2, DPAW08-000725-1, DoEE E2017-01290, and Fisheries collection exemption 2527 from the Department of Primary Industries and Regional Development) and the New Caledonia Province Sud (3252-2017/ARR/DENV).

## Acknowledgements

We thank the institutions that loaned us their valuable tissue samples, including the Museum and Art Gallery of the Northern Territory (MAGNT), South Australian Museum (SAM) and Western Australian Museum (WAM). We are also grateful to Isaac Rossetto, Tom Woods and Arne Rasmussen for assistance in collecting sea snake tissue samples. Finally, we thank Tessa Bradford for her guidance and support in the laboratory.

