## Supplementary Figures for "Seascape genomics reveals shared spatial genetic structure in three co-occurring sea snakes"

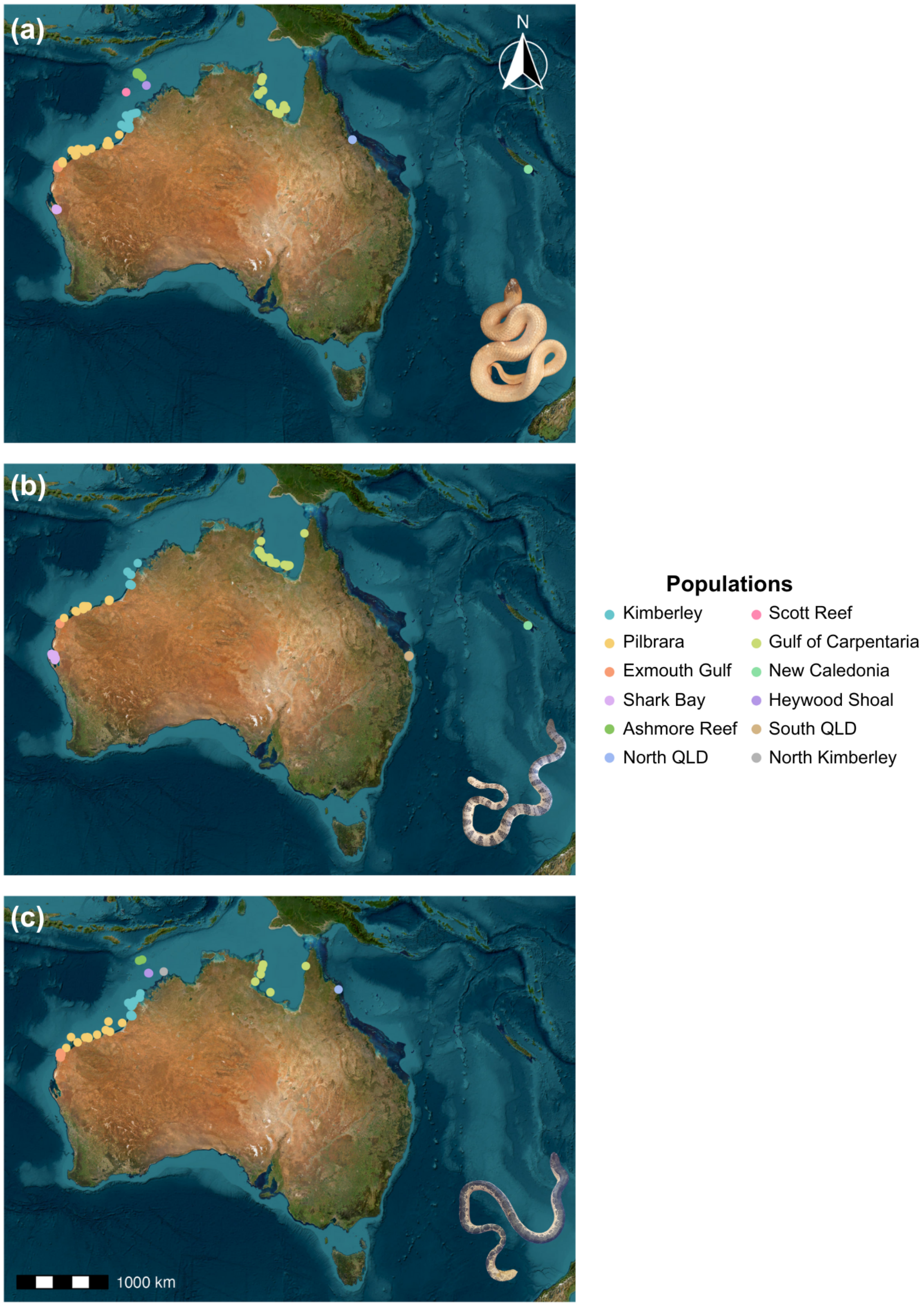


Figure S1: Map of sampling locations for (a) *Aipysurus laevis*, (b) *Hydrophis major*, and (c) *Hydrophis stokesii*. The ranges and population densities of each species varied between sampling regions, so not every species was sampled at each site. Specific information regarding the number of specimens collected at each site and other geographical information such as longitude and latitude can be found in Table S1. Colours of points represent regions. Photos of the three sea snakes are included (photographs taken by James Nankivell).

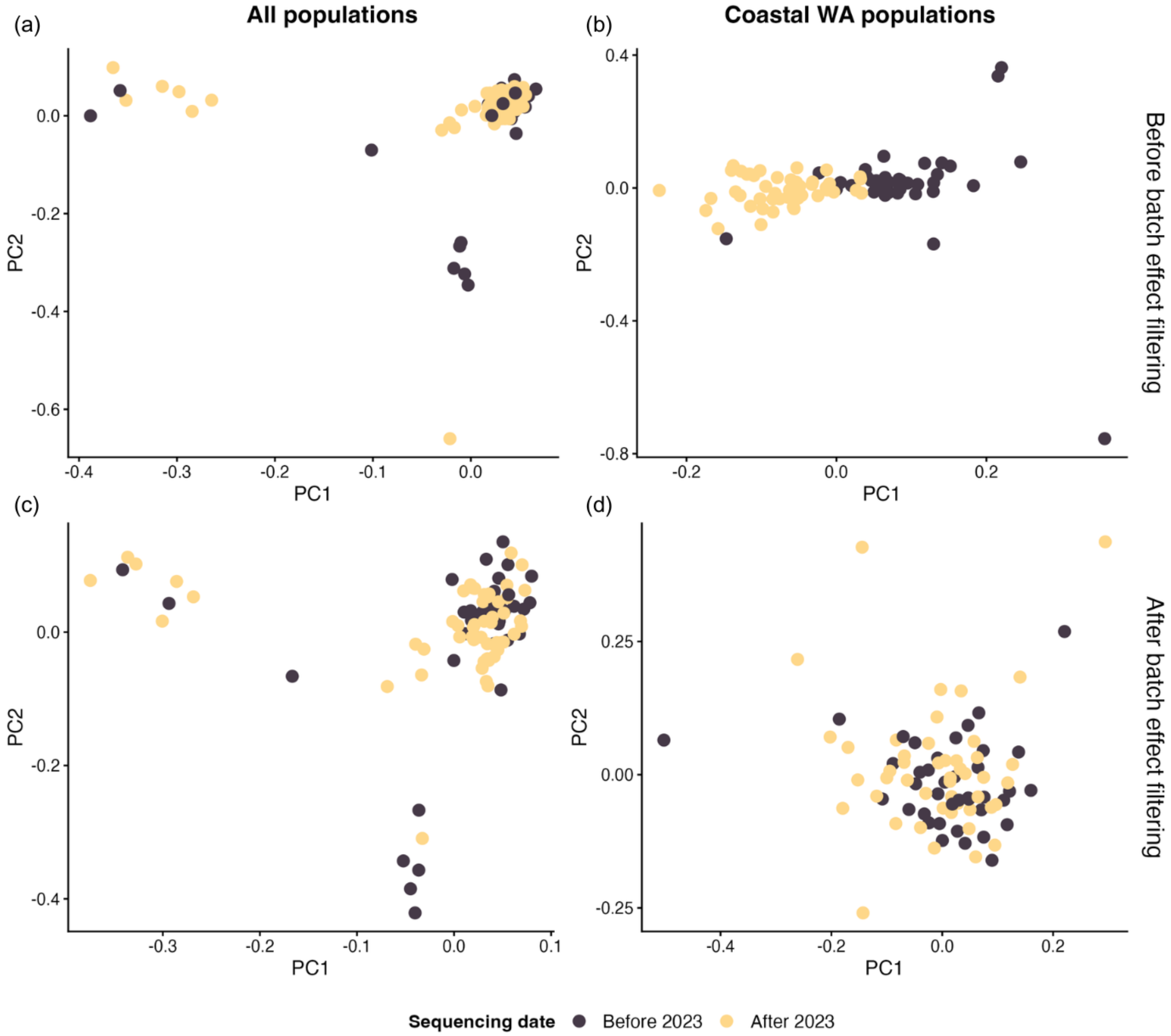


Figure S2: Principal Components Analyses (PCAs) of genetic data before (a, b) and after (c, d) removal of SNPs with absolute loadings ≥ 0.04 on PC1 of the coastal Western Australia (WA) PCA, used to identify batch effects. Panels show all populations (a, c) and coastal WA populations only (b, d).

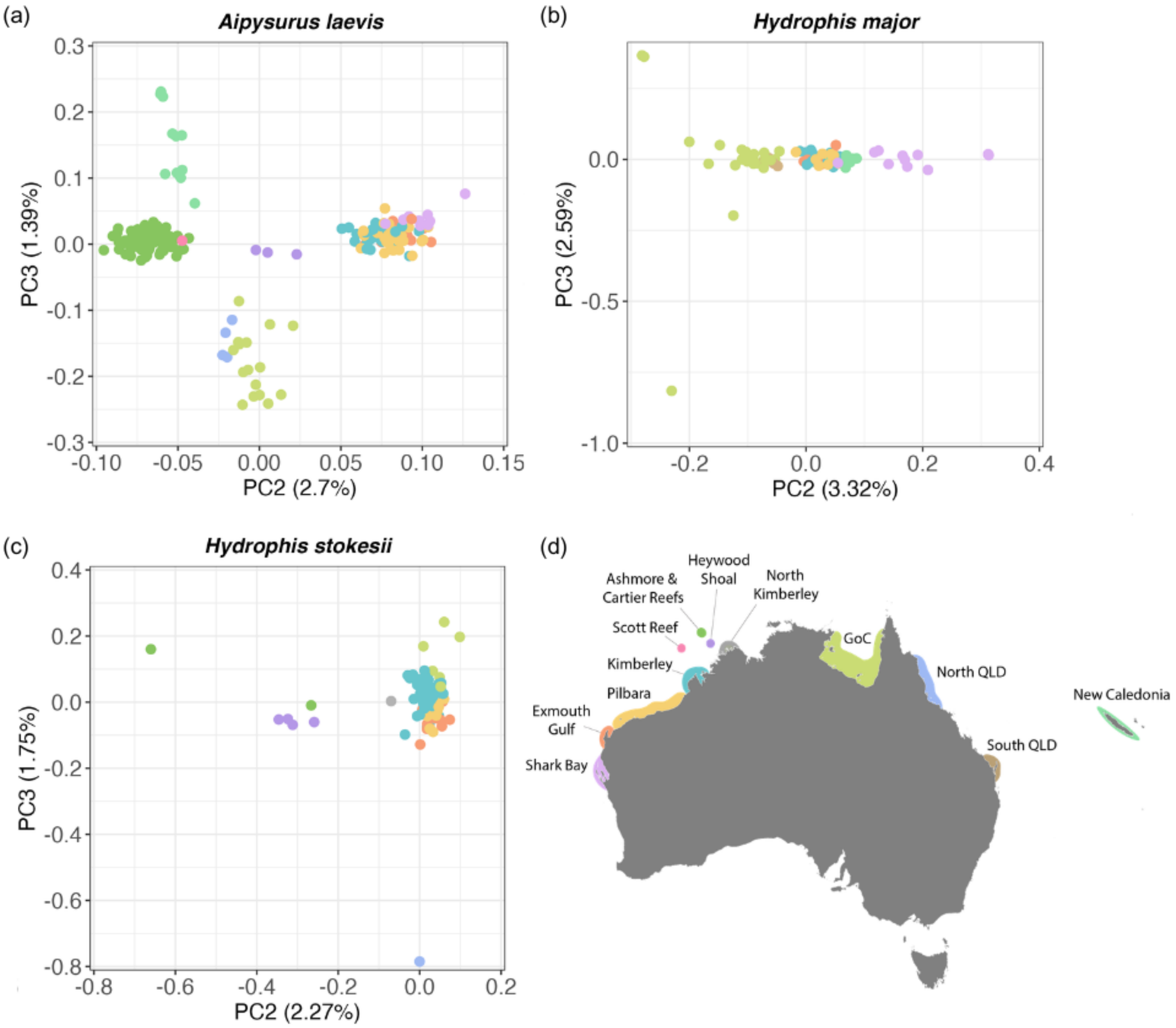


Figure S3: Principal Components (PCs) 2 and 3 of the Principal Components Analyses (PCA) performed on the genetic data from (a) *Aipysurus laevis*, (b) *Hydrophis major*, and (c) *Hydrophis stokesii* sampled across their Australian and New Caledonian ranges. Points are coloured according to the region of their origin as indicated in the legend map (d). The values indicated along each axis correspond to the percentage of inertia explained by the corresponding axis.

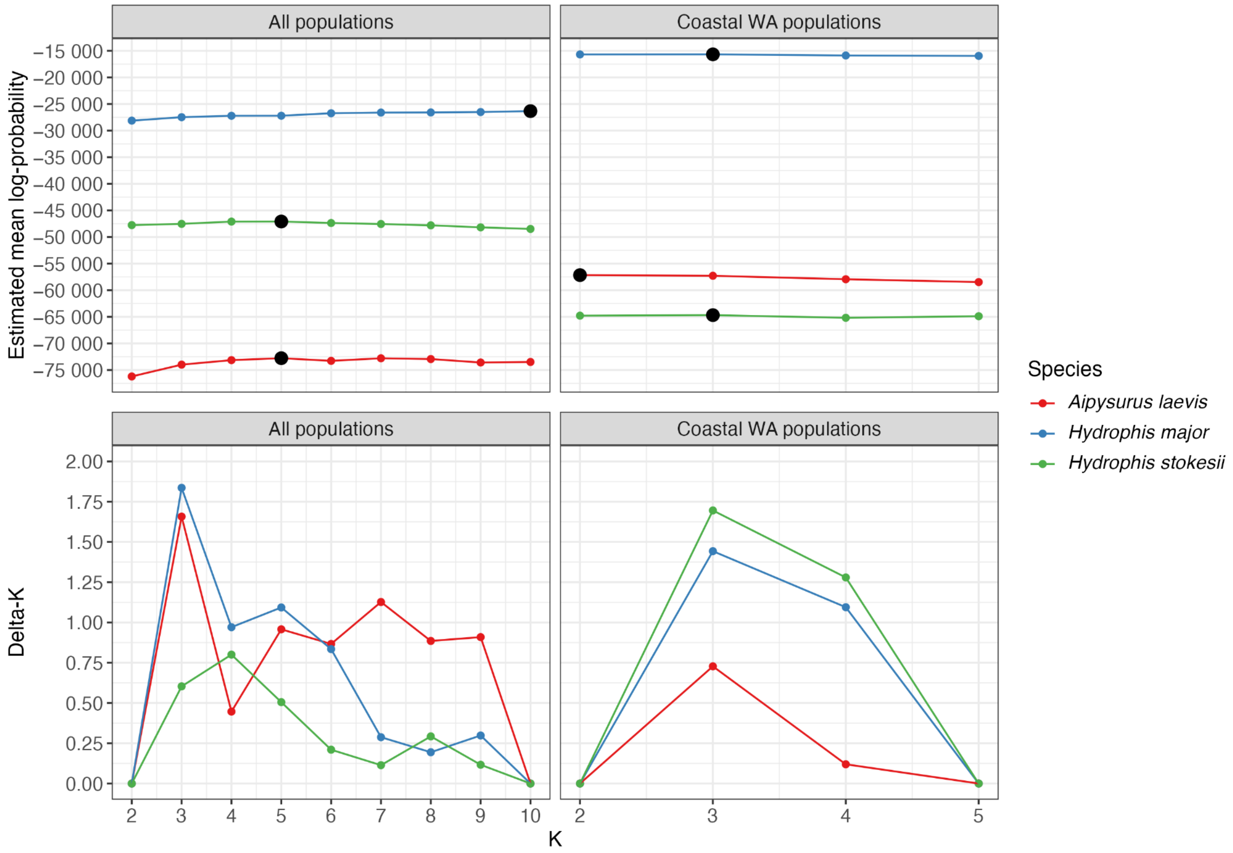


Figure S4: Estimated mean log-probability (top) and delta-*K* (Δ*K*) (bottom) for broad-scale (all populations) and regional-scale (coastal WA populations) STRUCTURE analyses, across different values of *K*. The mean log-probability plot shows the likelihood of the data given different numbers of genetic clusters (*K*), with higher values indicating better model support. The delta-*K* plot, calculated following Evanno et al., (2005), identifies the most likely number of genetic clusters by assessing the rate of change in log-probability between successive *K* values. Peaks in the delta-*K* plot indicate the optimal *K* values representing the most strongly supported population structure.

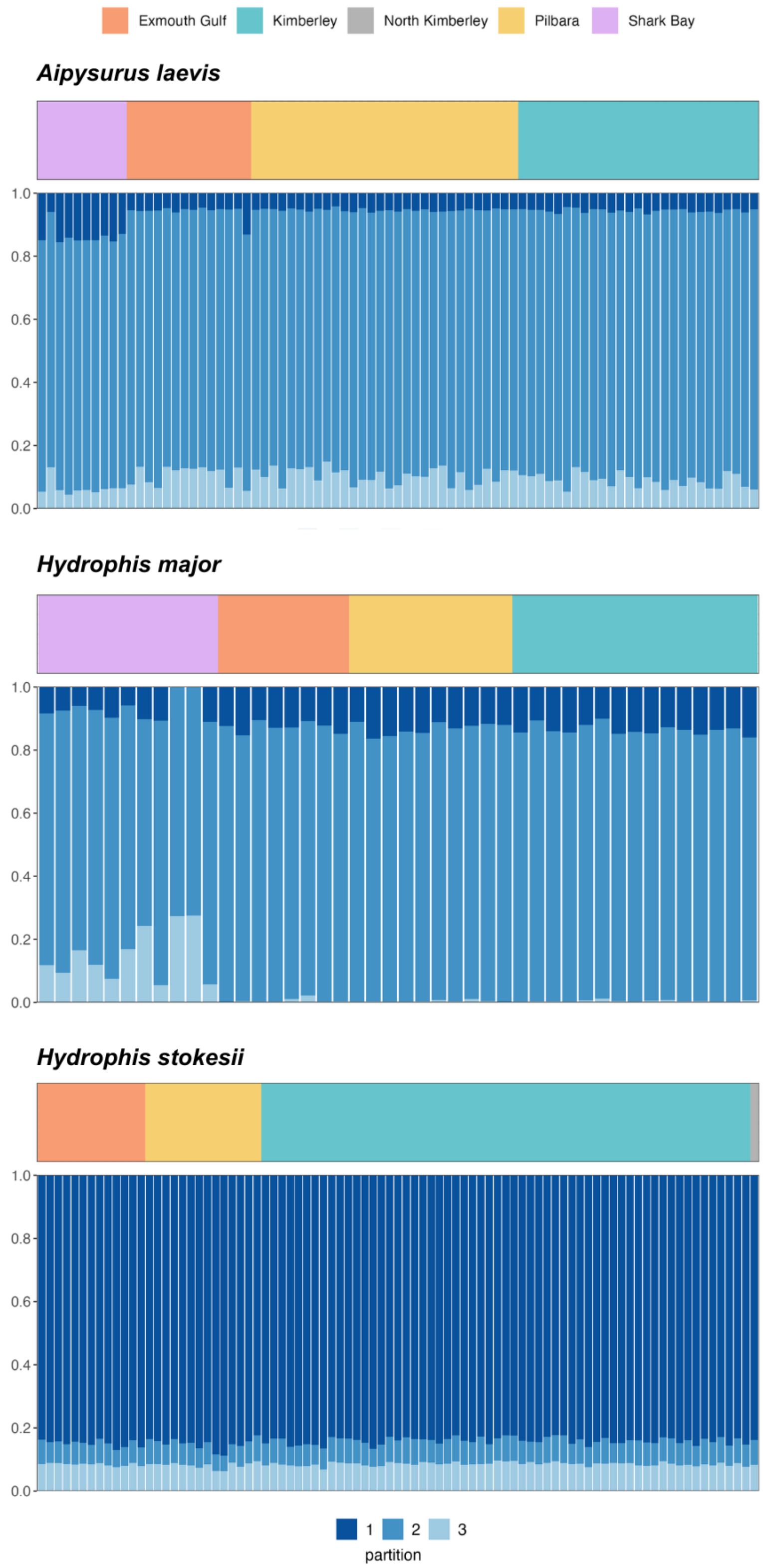


Figure S5: Ancestry profiles from regional-scale STRUCTURE analyses of coastal Western Australian populations, with region of origin indicated by colour in the top panel. Regions for both STRUCTURE and PCA plots are indicated in the legend map of Figure 3b.
